# Prioritizing the Valorization Strategies of an Invasive Fern (Azolla) in a Wetland

**DOI:** 10.1101/2024.03.07.583895

**Authors:** Farima Nikkhah, Mohammad Rahim Ramezanian, Kurt A. Rosentrater

## Abstract

Wetlands play a vital role as one of the most important natural habitats on our planet. However, the survival of these natural wetlands is threatened by various factors. The arrival of invasive and non-native aquatic ferns is one of these challenges. In this regard, Azolla filiculoides has become a severe problem for the Anzali wetland. Azolla, as an aquatic fern, has created numerous issues in aquatic habitats and paddy fields in recent decades. However, the valorization of Azolla can contribute to the establishment of a collection system for this invasive fern, which can consequently reduce the negative impact of this fern on the wetland, and it can serve as a free and available source of biomass. In this respect, a fuzzy multi-criteria decision-making approach was used to rank the valorization strategies of this invasive fern. Initially, through an in-depth literature review and expert opinions, four criteria were designated as indicators for research evaluation: 1) technical, 2) economic, 3) social, and 4) environmental. Six management options for Azolla were considered: 1) no collection, 2) collection and landfilling, 3) direct use as livestock and poultry feed, 4) composting, 5) biogas generation, and 6) biodiesel generation. The results revealed that “biodiesel generation,” “biogas generation,” and “composting” were ranked as the most effective management strategies for Azolla in the investigated wetland. This study suggests that bioenergy generation and compost production from Azolla are promising strategies towards mitigating the negative impact of this fern on the Anzali wetland.

## 1. Introduction

The world we live in today is vastly different from that of a century ago. We now inhabit a world that is densely populated and demands significant resources, equipped with advanced technologies that were completely absent in the last century. Environmental issues rank among the concerns that have consistently occupied individuals, governments, nations, and the entire world. In this context, wetlands serve an essential function as some of the most critical natural habitats on Earth. Natural wetlands are among the most active and vital ecosystems of the planet. Indeed, it is the wetlands that support the survival of certain animal and plant species. Wetlands are valuable assets currently threatened by various factors. The arrival of invasive and non-native aquatic ferns is one of these cases (Sadeghi et al., 2014). Non-native species do not naturally exist in one country, having evolved in another. The geographical range of these species is limited, and many of them are not naturally capable of crossing geographical barriers. However, humans have disrupted this pattern by transporting species across the globe. If such a species can survive, reproduce, and spread, it can be described as a biological invader. These species may damage biodiversity or negatively impact the operation of natural ecosystems. Currently, invasive species represent a significant threat to biodiversity and lead to considerable economic losses. Therefore, before the presence and spread of these species, coherent management measures must be taken to prevent widespread problems on biodiversity and human societies. A non-native species becomes invasive only when it spreads in a new area after settling in it, such that its distribution affects the environmental conditions of the site (Mardani and Ravanbakhsh, 2018). Certain conditions are required for a non-native species to become invasive. Azolla stands out as an interesting case because it is among the most invasive species globally, according to Pereira (2017). Over the past few decades, the Azolla aquatic fern has caused numerous issues in aquatic habitats. Azolla is a floating fern that can usually be observed in the stagnant and calm waters of wetlands and paddy fields in tropical areas. The genus Azolla belongs to the Pteridophyta genus, class Polypodiopsida, and order Salvinales, and has two subspecies and six living species (Roy et al., 2016). One of the unique characteristics of this fern is that it can quite easily adapt to new environments. This fern, with its thin roots, acts like a carpet that spreads across the water’s surface. Shortly after establishment, it occupies the entire surface, preventing sunlight from penetrating the depths. This lack of light penetration into the underwater layers halts or reduces the growth of submerged plants, leading to changes in the aquatic ecosystem. These changes present a major environmental challenge (Fallah et al., 2020).

Azolla, originally not from Iran, was brought into the country and quickly adapted to the favorable environmental conditions, resulting in its rapid spread. In just around forty years, it has become an invasive species, impacting aquatic ecosystems in the northern Iran. Initially introduced as a nitrogen-fixing fern, Azolla swiftly spread to the Anzali wetland and rice farms nearby. It was introduced for use as feed for livestock and poultry and as green manure. Azolla is now a major environmental concern for Anzali wetland. The presence of Azolla in aquatic environments, particularly in rice farms and the Anzali Wetland, has resulted in numerous problems, affecting these ecosystems (Sadeghi et al., 2012). The Anzali wetland is one of the valuable ecosystems in the Middle East (Jafari, 2009). Various solutions have been proposed to address the issues regarding the presence of Azolla in the region, but these efforts have not had significant results. However, Azolla offers several applications, including its use as a nitrogen fixer in rice farms, compost and plant fertilizers, animal feed, human nutrition, biomass energy resource, absorption and removal of phenols from wastewater, absorption of heavy metals from hydroponic systems, and biomonitoring of heavy metals from wastewater (Figure 1) (Hassanzadeh et al., 2021). It can be considered as a valuable source of protein, essential minerals, and vitamins, making it suitable for tropical climates and as livestock feed. Moreover, due to its relative high protein, carbohydrate, and crude fat content, concentrated Azolla protein has the potential to be used as a dietary supplement for humans, providing a gross energy content of 434.67 kcal per 100 g (Mohammad et al., 2018).

**Figure 1.**
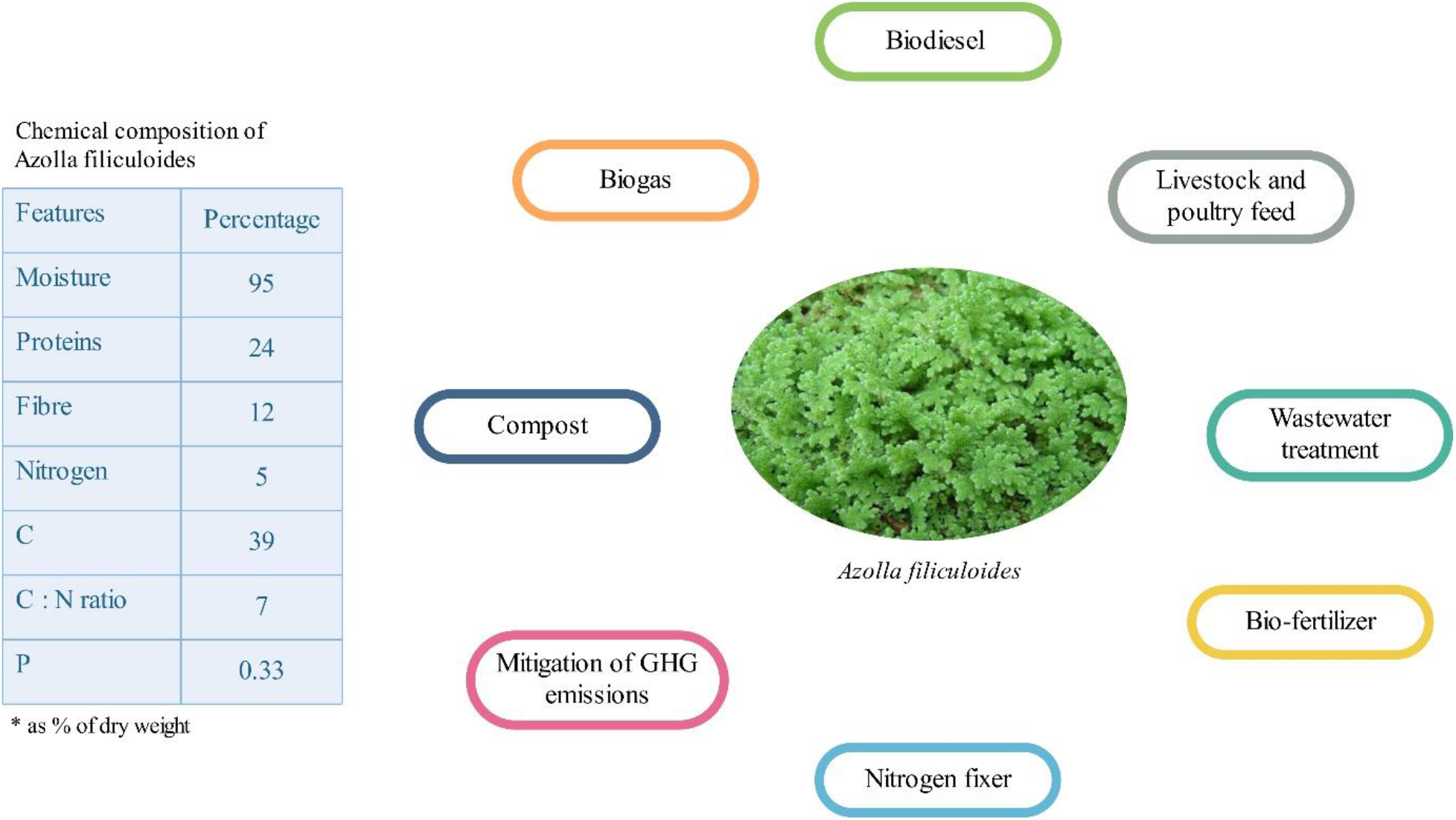
Characterization and application of Azolla filiculoides (Singh et al., 1981; Kamaruddin et al., 2019)

Considering the aforementioned applications of Azolla, it is importnat to prioritize the valorization strategies of this fern in wetlands. The establishment of a collection system for this invasive species not only helps mitigate its negative impact on the wetland but also allows for the utilization of Azolla as an easily accessible biomass resource. In this study, a fuzzy multi-criteria decision-making method was used to establish the ranking of Azolla valorization strategies mentioned in Figure 1 in the Anzali Wetland.

## 2. Material and methods

### 2.1. Study area

The Anzali Wetland was chosen as the case study. It is located in the northern region of Iran, within the Guilan province, near the port of Anzali. This wetland, spanning about 20,000 hectares, lies between saline and freshwater sources in the southwest of the Caspian Sea, adjacent to the Sefidrood Delta. Figure 2 shows the location of the Anzali wetland.

**Figure 2.**
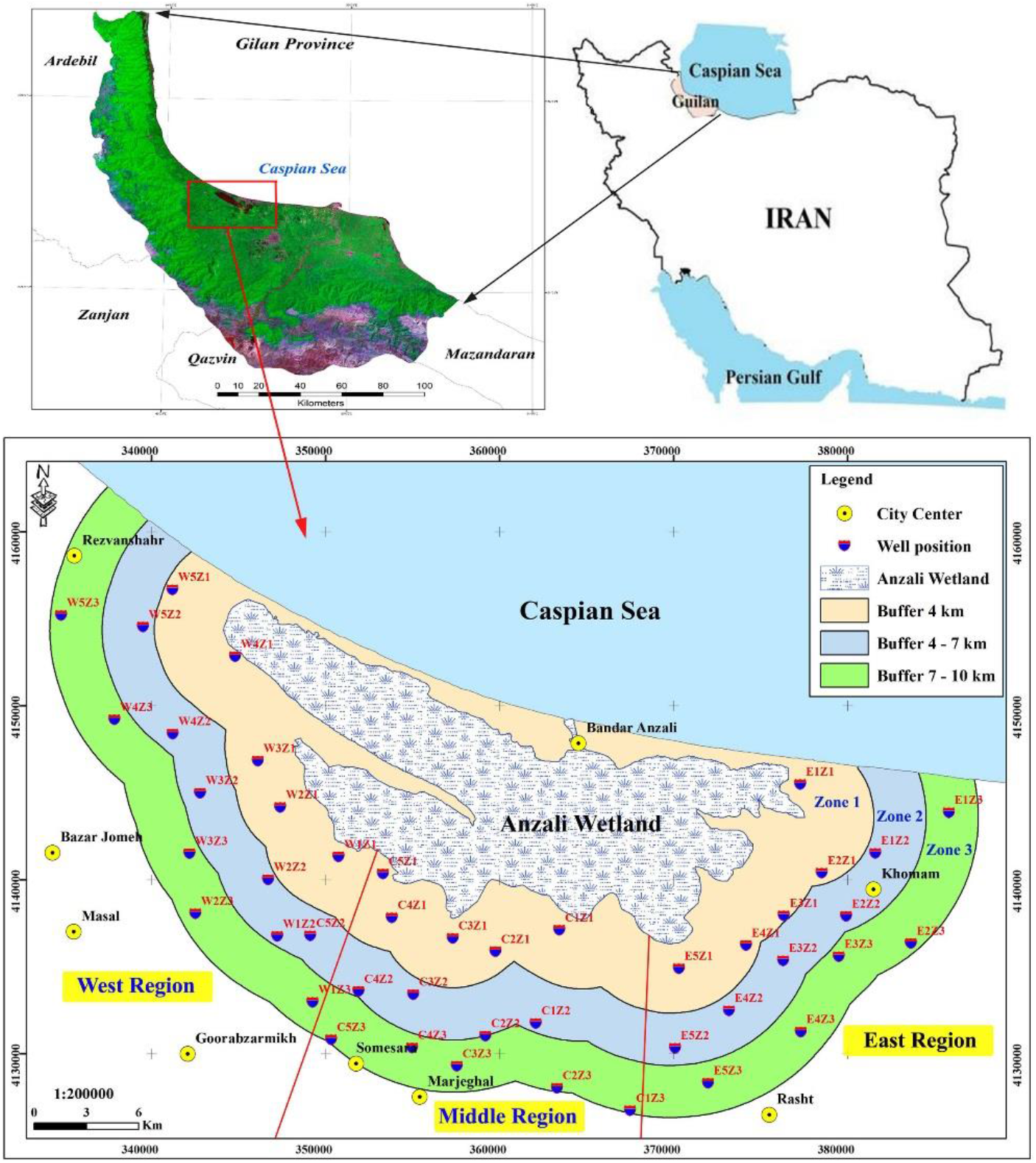
Location of Anzali wetland (Vatandoost et al., 2018)

### 2.2 Decision-making tree

In this study, a practical research approach was adopted, involving the collection of data through a structured questionnaire. To prioritize the valorization strategies for Azolla in the Anzali Wetland, a fuzzy-based multi-criteria decision-making approach was applied. It is important to identify criteria to evaluate the options when dealing with a fuzzy multi-criteria decision-making problem. The questionnaire was designed following a literature review and consultations with experts in the field. The four decision criteria were: 1) technical, 2) economic, 3) social, and 4) environmental (Table 1).

**Table 1.** Criteria considered to prioritize the Azolla applications.

| Code | Criteria | Row |
| --- | --- | --- |
| C1 | Technical | 1 |
| C2 | Economic | 2 |
| C3 | Social | 3 |
| C4 | Environmental | 4 |

This study listed six strategies for Azolla management in Anzali wetland: 1) no collection, 2) collection and landfilling, 3) direct use as livestock and poultry feed, 4) composting, 5) biogas generation, and 6) biodiesel generation (Table 2).

**Table 2.** Valorization strategies considered to prioritize the Azolla applications.

| Code | Options | Row |
| --- | --- | --- |
| A1 | No collection | 1 |
| A2 | Collection and landfill | 2 |
| A3 | Direct use as livestock and poultry feed | 3 |
| A4 | Composting | 4 |
| A5 | Biogas generation | 5 |
| A6 | Biodiesel generation | 6 |

In the next step, based on the considered criteria and options, the hierarchical tree of the research is categorized into three levels: the first level (the target level) focuses on the prioritization of Azolla management methods; the second level is the criteria level, which includes technical, economic, social, and environmental factors; and the third level is the options level, which encompasses no collection, collection and landfilling, direct use as livestock and poultry feed, composting, biogas generation, and biodiesel generation (Figure 3).

**Figure 3.**
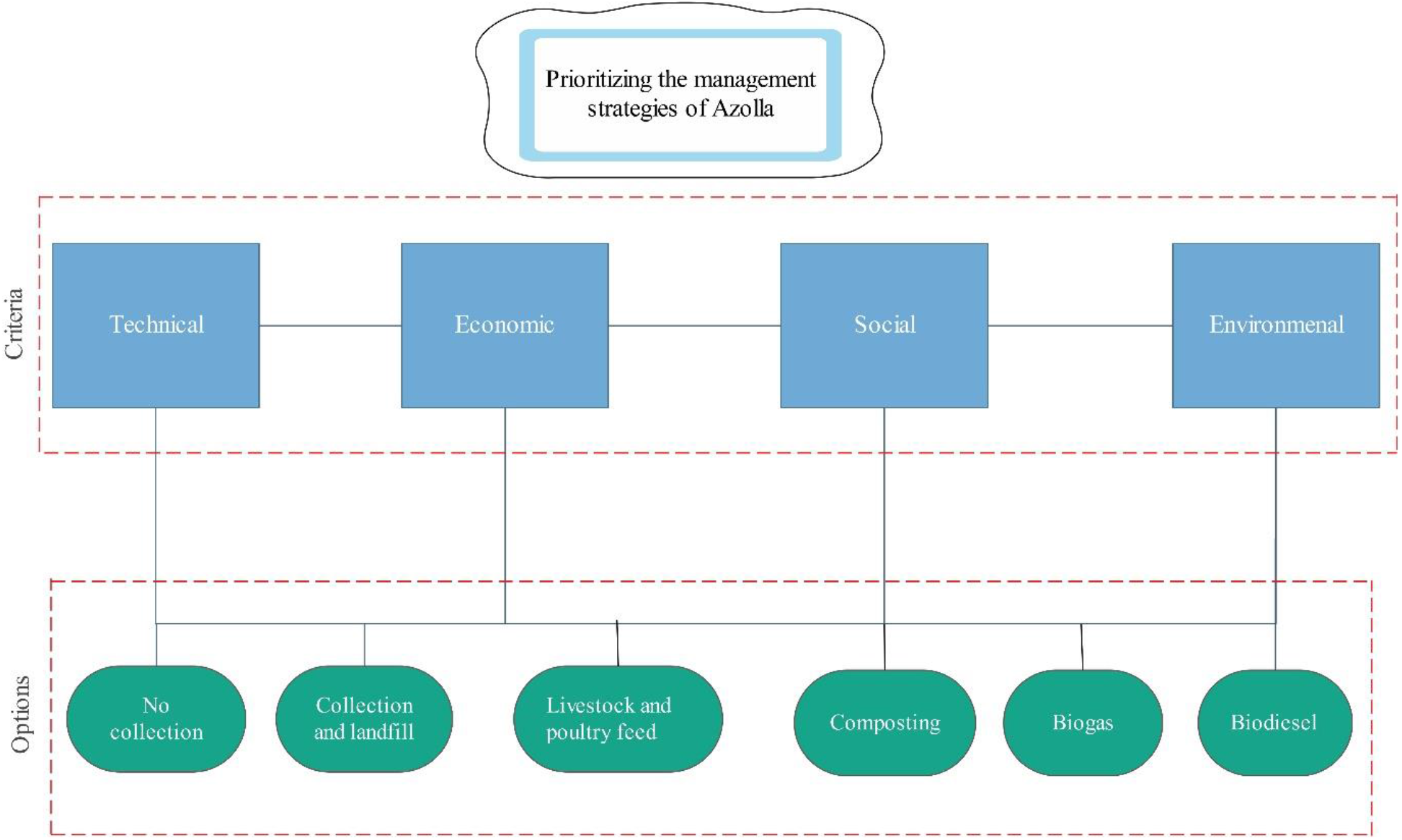
Decision making tree of determining the best option of Azolla management

A questionnaire was distributed to 12 experts in the field of study. Initially, the criteria were compared pairwise; for example, the economic criterion was compared to the social criterion. Subsequently, the options were compared in pairs considering each criterion. The experts were asked to use a score scale of 1 to 9 for the pairwise comparisons. For example, if an expert assigns a score of 3 to the comparison of technical versus economic criteria, then, according to Table 3, the triangular fuzzy number 3 is represented as (L, M, U) = (2, 3, 4). If this preference is -3, it equals (1/U, 1/M, 1/L), which is (1/4, 1/3, 1/2).

**Table 3.**
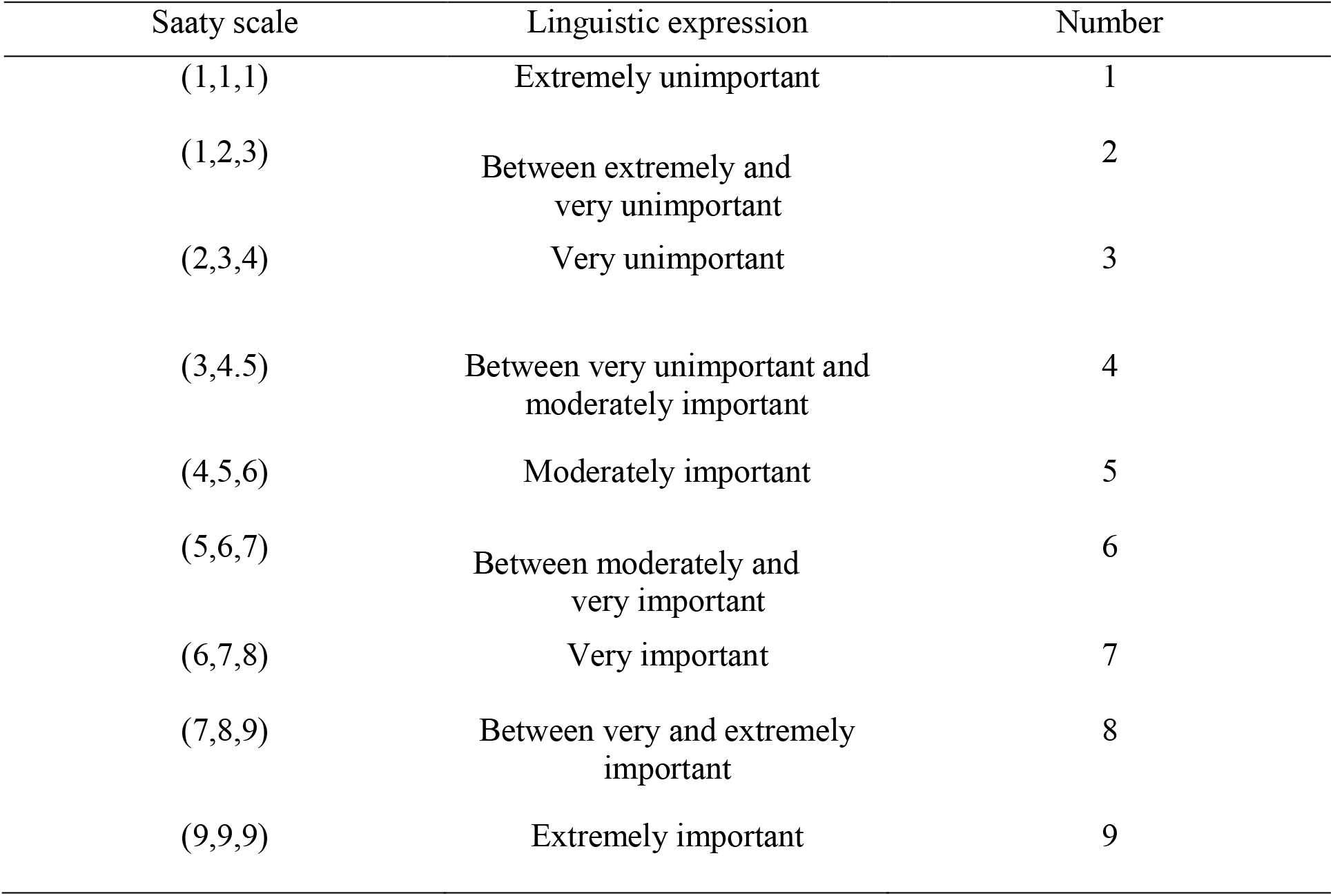
Saaty scale and the corresponding triangular fuzzy numbers (Kaganski et al, 2018)

| Saaty scale | Linguistic expression | Number |
| --- | --- | --- |
| (1,1,1) | Extremely unimportant | 1 |
| (1,2,3) | Between extremely and very unimportant | 2 |
| (2,3,4) | Very unimportant | 3 |
| (3,4,5) | Between very unimportant and moderately important | 4 |
| (4,5,6) | Moderately important | 5 |
| (5,6,7) | Between moderately and very important | 6 |
| (6,7,8) | Very important | 7 |
| (7,8,9) | Between very and extremely important | 8 |
| (9,9,9) | Extremely important | 9 |

### 2.3 Fuzzy multi-criteria decision making

The Analytic Hierarchy Process (AHP), developed by Saaty in 1980, was used as the multi-criteria decision making method. The AHP technique is a multi-criteria decision-making that assesses the weights of criteria and prioritizes options in a structured manner through pairwise comparisons (Bao et al., 2021). Considering that linguistic values and subjective judgments during comparisons can be imprecise, this study utilized fuzzy logic (Maceika et al., 2021). Buckley (1985) expanded Saaty’s AHP by assigning precise ratios when comparing criteria and options, and by deriving their fuzzy weights using the geometric mean method. In this study, triangular fuzzy numbers and the 9-digit spectrum (Table 3) were used.

As shown in Table 3, fuzzy numbers are defined by their lower (L), middle (M), and upper (U) limits: L refers the lower bounds of the fuzzy number, M to the middle limit, and U to the upper bounds. Equation (1) displays the Triangular Fuzzy Number (TFN) membership function (Azar and Rajabzade, 2017). A TFN should exhibit the following basic features, as depicted by Equations (1-6) (Laarhoven and Pedrycz, 1983):

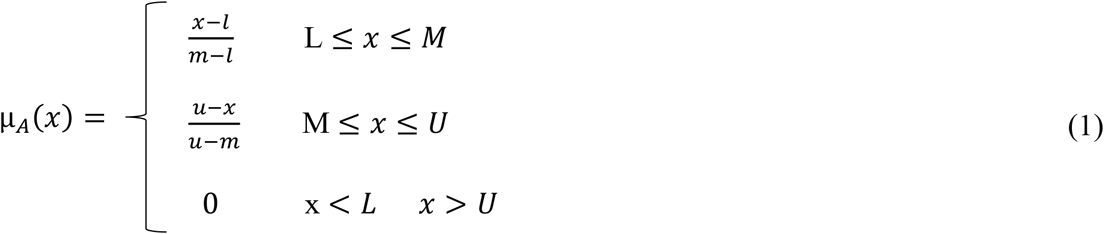

Figure 4 illustrates the membership function of the triangular fuzzy number, L and U represent the lower and upper bounds of the fuzzy number *Ā*, respectively, and M is the modal value. TFN can be shown by *Ā* = (*L. M. U*) and the operational laws of two TFNs *Ā*_1_ =(L_1_,M_1_,U_1_) and Ã_2_ =(L_2_,M_2_,U_2_) are explained as follows (Hsieh et al., 2004):

**Figure 4.**
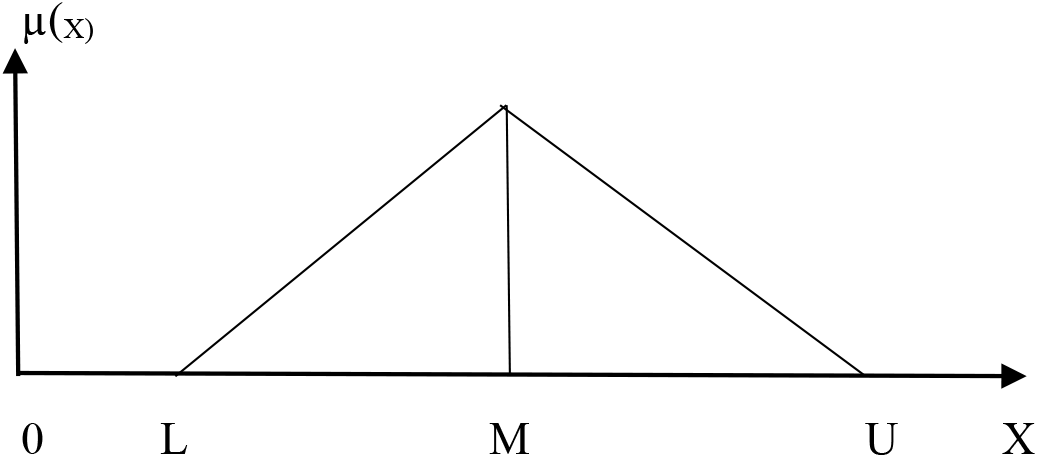
Membership function of the triangular fuzzy number (Azar and Rajabzade, 2017)

Addition of two fuzzy numbers: (+)

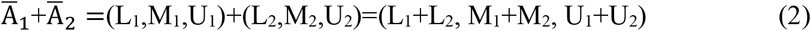

Multiplication of two fuzzy numbers: (×)

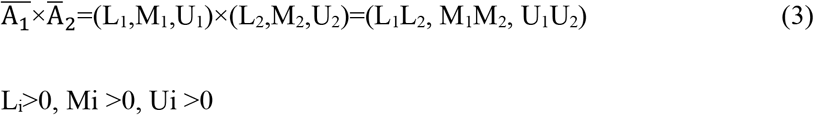

Subtraction of two fuzzy numbers: (-)

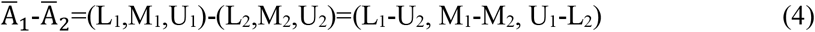

Division of two fuzzy numbers: (÷)

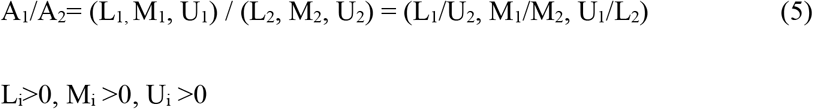

Reciprocal of a fuzzy number

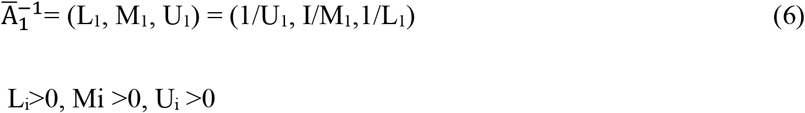

In an Excel environment, a 4×4 square matrix was initially formed, where the rows and columns represented the criteria, with each cell indicating a paired comparison. The primary diagonal numbers were set to one, and the numbers above and below the main diagonal were inverses of each other, as demonstrated in Equation 7:

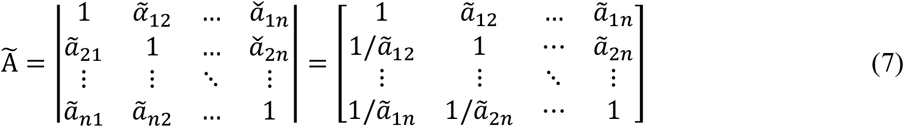

In this way, it consists of four 6x6 matrices where the rows and columns represent options filled with fuzzy numbers. In the next step, the comparisons were consolidated into pairwise comparisons, and the geometric means of the rows were calculated. First, the geometric mean of the lower limit (L), followed by the geometric mean of the middle limit (M), and then the upper limit (U) were calculated using the Buckley geometric mean method (Equation 8). Subsequently, the geometric means of each column were summed, and each geometric mean was multiplied by the inverse of this sum as per Equation 9. The fuzzy weights (W) were converted to defuzzified values using Equation 10. Finally, the normalized weights of the criteria relative to the target and the options relative to the criteria were computed by dividing the defuzzified weight by the total weights. After obtaining the normalized weights with the Buckley method, the final weight was determined using the SAW method.

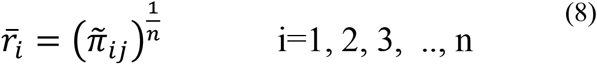

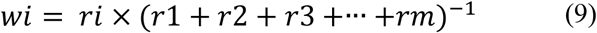

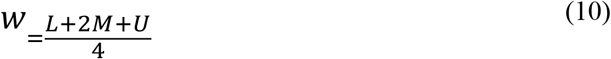

where n is the number of related elements in each row, and 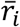 is the geometric mean of the fuzzy comparison value of criterion i to each other criterion. In addition, incompatibility rates were calculated after the comparisons were consolidated.

### Fuzzy-based AHP consistency verification

Multiply each entry in column i of matrix Ã by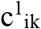, then divide the summation of values in row i by 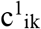 to yield another column vector, where 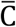 refers to a weighted sum vector. Then, compute the averages of values in vector 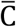 to obtain the maximum eigenvalue of matrix Ã, as shown in equations 11,12, and 13 (Ho et al, 2012).

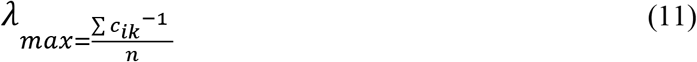

Compute the consistency index,

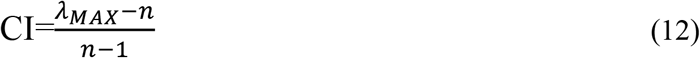

Compute the consistency ratio,

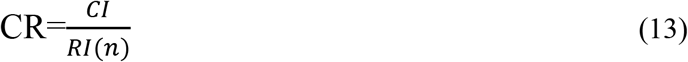

Where RI(n) is a random index whose value depends on the value of n, that Table 4 presents the incompatibility rates of the merged matrix. The incompatibility rate should always be less than 0.1 for the comparisons to be deemed appropriately consistent. If this rate exceeds 0.1, the comparisons need to be revised. After examining the incompatibility rates of the pairwise comparisons made by the experts, all were found to be consistent.

**Table 4.**
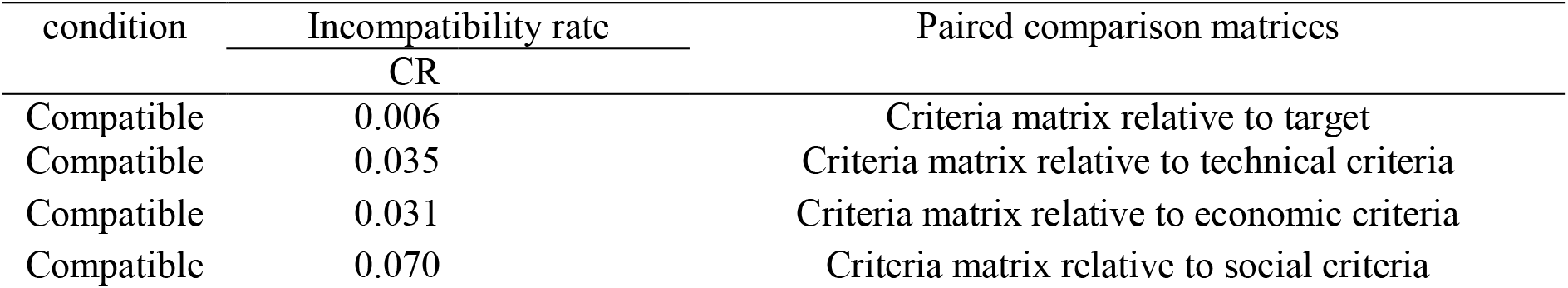
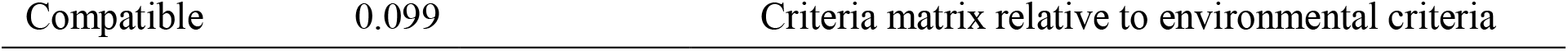
Incompatibility rates of the merged matrix.

### Simple Additive Weighting

The options were prioritized, and a decision matrix (agreement table) was prepared, with criteria as columns and options as rows. The normalized weight of each valorization strategy was then entered in the corresponding column under each criterion. The weight of the criteria, resulting from their pairwise comparison with the target, was also placed in the bottom row. The logic of the Simple Additive Weighting (SAW) method was employed to determine the weight of the alternatives, and the weight of each alternative was obtained using this decision-making method. It should be noted that the limited number of available experts in the studied area constituted one of the limitations of the current study. Furthermore, only a limited number of criteria/options were considered in this study to conserve the experts’ energy and time by not requiring them to consider less important criteria/options.

## 3. Results and discussion

### 3.1. Prioritization of criteria

Table 5 displays the fuzzy, definite, and normalized weights of the criteria. The results of the criteria ranking indicated that the economic criteria, with a normalized weight of 0.35, had the highest priority in the valorization of Azolla in the Anzali Wetland. The environmental criteria, with a normalized weight of 0.25, were ranked as the second priority. Technical criteria, with a normalized weight of 0.21, held the third priority. Social criteria, with a normalized weight of 0.20, had the lowest priority.

**Table 5.** Weights of the selected criteria in the valorization of Azolla in the Anzali wetland.

| Criteria | Normalized weights | Definite weights | Fuzzy weights criteria |  |  | Geometric mean of each row |  |  | Rank |
| --- | --- | --- | --- | --- | --- | --- | --- | --- | --- |
|  |  |  | U | M | L | U | M | L |  |
| C1 | 0.21 | 0.21 | 0.29 | 0.21 | 0.15 | 0.90 | 0.76 | 0.65 | 3 |
| C2 | 0.35 | 0.36 | 0.50 | 0.34 | 0.24 | 1.56 | 1.26 | 1.05 | 1 |
| C3 | 0.20 | 0.20 | 0.28 | 0.20 | 0.14 | 0.88 | 0.72 | 0.61 | 4 |
| C4 | 0.25 | 0.26 | 0.36 | 0.25 | 0.18 | 1.11 | 0.91 | 0.79 | 2 |
| Sum | 1.00 | 1.03 |  |  |  | 4.44 | 3.66 | 3.10 |  |

### 3.2. Prioritization of valorization strategies

#### 3.2.1. Prioritization based on the technical criteria

Table 6 presents the fuzzy, definite, and normalized weights of the options according to the technical aspect of valorization strategies. Accordingly, the generation of biogas, with a normalized weight of 0.28, had the highest priority. The composting strategy, with a normalized weight of 0.24, was placed in the second priority. The generation of biodiesel, with a normalized weight of 0.21, occupied the third priority. Direct usage of Azolla as livestock and poultry feed, with a normalized weight of 0.12, was the fourth priority. Collection and landfilling of Azolla, with a weight of 0.09, was identified as the fifth priority. Non-collection of Azolla was placed in the last priority.

**Table 6.** Weights of the options in the light of the technical criteria.

| Technical-based prioritization | Definite weights | Fuzzy weights |  |  | Geometric average of rows |  |  | Normalized weight | Rank |
| --- | --- | --- | --- | --- | --- | --- | --- | --- | --- |
|  |  | U | M | L | U | M | L |  |  |
| A1 | 0.06 | 0.09 | 0.06 | 0.04 | 0.73 | 0.60 | 0.49 | 0.06 | 6 |
| A2 | 0.10 | 0.14 | 0.09 | 0.06 | 1.21 | 0.99 | 0.81 | 0.09 | 5 |
| A3 | 0.13 | 0.19 | 0.12 | 0.08 | 1.58 | 1.34 | 1.11 | 0.12 | 4 |
| A4 | 0.25 | 0.37 | 0.24 | 0.15 | 3.09 | 2.52 | 1.98 | 0.24 | 2 |
| A5 | 0.29 | 0.43 | 0.28 | 0.18 | 3.62 | 2.99 | 2.31 | 0.28 | 1 |
| A6 | 0.22 | 0.34 | 0.21 | 0.13 | 2.83 | 2.26 | 1.72 | 0.21 | 3 |
| Sum | 1.05 |  |  |  | 13.06 | 10.71 | 8.41 | 1.00 |  |

#### 3.2.2. Prioritization based on the economic criteria

The fuzzy, definite, and normalized weights of the valorization options based on the technical criteria are displayed in Table 7. The generation of biodiesel, with a normalized weight of 0.32, had the highest priority. Biogas generation was ranked as the second priority based on economic criteria. Composting, with a normalized weight of 0.20, was in the third priority. The direct usage of Azolla as livestock and poultry feed, with a normalized weight of 0.13, was the fourth priority. Collection and landfilling of Azolla were placed in the fifth priority.

**Table 7.** Weights of options relative to economic criteria.

| Economic-based prioritization | Normalized weights | Definite weights | Fuzzy weights |  |  | Geometric average of rows |  |  | Rank |
| --- | --- | --- | --- | --- | --- | --- | --- | --- | --- |
|  |  |  | U | M | L | U | M | L |  |
| A1 | 0.03 | 0.04 | 0.05 | 0.03 | 0.02 | 0.41 | 0.33 | 0.28 | 6 |
| A2 | 0.04 | 0.05 | 0.07 | 0.04 | 0.03 | 0.52 | 0.41 | 0.33 | 5 |
| A3 | 0.13 | 0.13 | 0.21 | 0.13 | 0.08 | 1.57 | 1.21 | 0.95 | 4 |
| A4 | 0.20 | 0.21 | 0.32 | 0.20 | 0.13 | 2.44 | 1.96 | 1.52 | 3 |
| A5 | 0.27 | 0.28 | 0.42 | 0.27 | 0.17 | 3.15 | 2.59 | 2.03 | 2 |
| A6 | 0.32 | 0.33 | 0.49 | 0.32 | 0.21 | 3.68 | 3.06 | 2.43 | 1 |
| Sum | 1.00 | 1.05 |  |  |  | 11.77 | 9.56 | 7.53 |  |

#### 3.2.3. Prioritization based on the social criteria

Table 8 displays the fuzzy, definite, and normalized weights of the options in light of social criteria. The generation of biodiesel, with a normalized weight of 0.39, had the highest priority. Biogas generation was placed in the second priority, with a normalized weight of 0.22. Composting, with a normalized weight of 0.19, was in the third priority. The fourth priority was the direct application of Azolla as feed for livestock and poultry. Collection and landfilling, with a weight of 0.05, was placed as fifth priority.

**Table 8.** Weights of the options in the case of social criteria.

| Social-based prioritization | Normalized weight | Definite weights | Fuzzy weights |  |  | Geometric average of row |  |  | Rank |
| --- | --- | --- | --- | --- | --- | --- | --- | --- | --- |
|  |  |  | U | M | L | U | M | L |  |
| A1 | 0.03 | 0.03 | 0.04 | 0.02 | 0.02 | 0.43 | 0.34 | 0.28 | 6 |
| A2 | 0.05 | 0.05 | 0.07 | 0.05 | 0.03 | 0.79 | 0.63 | 0.52 | 5 |
| A3 | 0.12 | 0.12 | 0.16 | 0.12 | 0.09 | 1.84 | 1.57 | 1.31 | 4 |
| A4 | 0.19 | 0.20 | 0.26 | 0.19 | 0.15 | 2.98 | 2.57 | 2.19 | 3 |
| A5 | 0.22 | 0.23 | 0.28 | 0.23 | 0.18 | 3.23 | 2.99 | 2.75 | 2 |
| A6 | 0.39 | 0.39 | 0.50 | 0.39 | 0.30 | 5.75 | 5.13 | 4.49 | 1 |
| Sum | 1.00 | 1.02 |  |  |  | 15.02 | 13.23 | 11.55 |  |

#### 3.2.4. Prioritization based on the environmental criteria

Generation of biogas, with a normalized weight of 0.30, was ranked as the highest priority based on the environmental criteria (Table 9). Valorization of Azolla through biodiesel generation was the second priority. Composting, with a normalized weight of 0.21, was the third priority.

**Table 9.** Weights of the options in the light of the environmental criteria.

| Environmental-based prioritization | Normalized weights | Definite weights | Fuzzy weights |  |  | Geometric average of rows |  |  | Rank |
| --- | --- | --- | --- | --- | --- | --- | --- | --- | --- |
|  |  |  | U | M | L | U | M | L |  |
| A1 | 0.02 | 0.03 | 0.03 | 0.02 | 0.02 | 0.33 | 0.29 | 0.25 | 6 |
| A2 | 0.06 | 0.06 | 0.08 | 0.05 | 0.04 | 0.77 | 0.63 | 0.54 | 5 |
| A3 | 0.12 | 0.12 | 0.16 | 0.12 | 0.09 | 1.64 | 1.37 | 1.16 | 4 |
| A4 | 0.21 | 0.22 | 0.29 | 0.21 | 0.16 | 2.85 | 2.47 | 2.12 | 3 |
| A5 | 0.30 | 0.30 | 0.39 | 0.30 | 0.22 | 3.90 | 3.45 | 2.98 | 1 |
| A6 | 0.29 | 0.29 | 0.37 | 0.29 | 0.22 | 3.70 | 3.31 | 2.88 | 2 |
| Sum | 1.00 | 1.02 |  |  |  | 13.20 | 11.53 | 9.93 |  |

The SAW method was applied to integrate the final ranks of the options, and the results are displayed in Table 10. Overall, the study’s results indicated that the generation of biodiesel, with a normalized weight of 0.30, had the highest priority. Biogas, with a normalized weight of 0.27, was identified as the second priority. Composting Azolla, with a normalized weight of 0.21, was the third priority. Using Azolla directly as feed for livestock and poultry, with a normalized weight of 0.12, was ranked as the fourth priority. Collection and landfilling, with a weight of 0.06, were placed fifth.

**Table 10.**
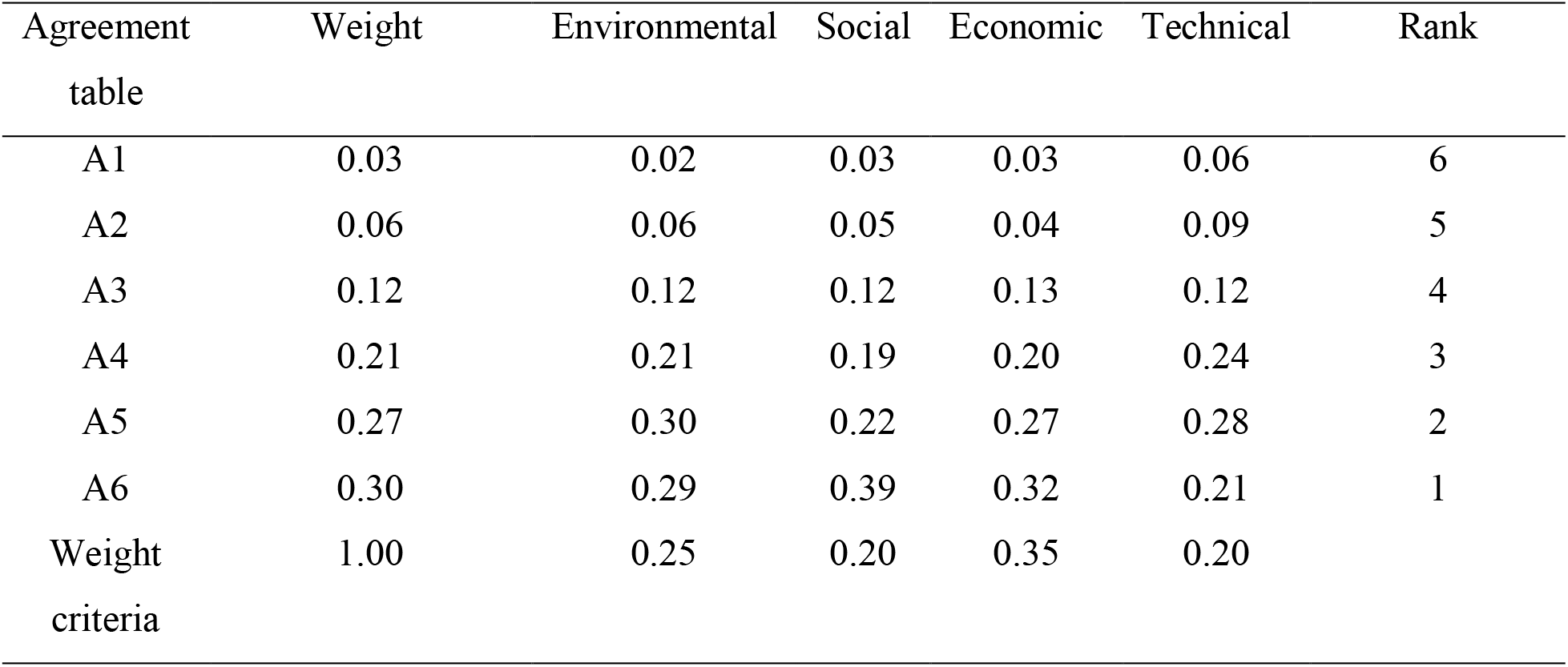
Agreement table (decision matrix)

| Agreement table | Weight | Environmental | Social | Economic | Technical | Rank |
| --- | --- | --- | --- | --- | --- | --- |
| A1 | 0.03 | 0.02 | 0.03 | 0.03 | 0.06 | 6 |
| A2 | 0.06 | 0.06 | 0.05 | 0.04 | 0.09 | 5 |
| A3 | 0.12 | 0.12 | 0.12 | 0.13 | 0.12 | 4 |
| A4 | 0.21 | 0.21 | 0.19 | 0.20 | 0.24 | 3 |
| A5 | 0.27 | 0.30 | 0.22 | 0.27 | 0.28 | 2 |
| A6 | 0.30 | 0.29 | 0.39 | 0.32 | 0.21 | 1 |
| Weight criteria | 1.00 | 0.25 | 0.20 | 0.35 | 0.20 |  |

Recent studies highlight that Azolla is a valuable source of biomass for biofuel production (Salehzadeh et al., 2014; Brouwer et al., 2016; Roy et al., 2016). For example, Arefin et al. (2021) showed that Azolla stands out as a promising fern for producing high-quality biofuels relative to other studied aquatic ferns. The studied country (Iran) ranks among the top 10 global emitters of greenhouse gas (GHG) emissions (Nikkhah, 2018; Firouzi et al., 2018). This country continues to heavily depend on the limited resources of non-renewable fossil fuels (Bastan and Shakouri, 2018; Rajabi et al., 2021). Generating a portion of the energy from biomass resources can help mitigate GHG emissions. In this context, Azolla, which is available in bulk quantities as a free and accessible resource in the northern regions of Iran, presents a sustainable option for biofuel production. Additionally, Azolla is considered as a valuable source of biomass for composting (Razavipour et al., 2018). Azolla and its compost can be effectively utilized to enhance the yield of agricultural products, such as rice (Thapa and Poundel, 2021) and maize (Maswada et al., 2021).

## 4. Conclusions and future directions

Azolla-an invasive fern-has occupied a large area of Anzali wetland. This fern with overgrowth and uncontrolled reproduction has become a major environmental challenge for the wetland, however Azolla has many applications. In this respect, the fuzzy analytic hierarchy process (fuzzy-AHP) was used to address this complex problem in order to prioritize the valorization strategies of Azolla. Four criteria and six valorization strategies were considered in this study. The results indicated that biofuel generation and composting are the best options for managing this invasive fern in Anzali. Further research is needed to provide insights into the sustainability (economic, environmental, and social aspects) of biofuel generation and composting Azolla in the studied area.

## Ethical procedure

As an expert scientist, along with co-authors from the concerned field, this paper has been submitted with full responsibility, adhering to the due ethical procedures. There is no duplicate publication, fraud, plagiarism, or concerns regarding animal or human experimentation.

## Credit authorship contribution statement

Farima Nikkhah contributed to developing the initial idea, data collection, analysis and writing the results and discussion section. Mohammad Rahim Ramezanian contributed to editing of the manuscript. Kurt Rosentrater contributed to the editing and supervising of the manuscript.

## References

Arefin, M.A., Rashid, F. and Islam, A., 2021. A review of biofuel production from floating aquatic plants: an emerging source of bio-renewable energy. Biofuels, Bioproducts and Biorefining, 15(2), 574–591.

Azar, A., Rajabzade, A. (2017). Multi-Attribute Decision Making(MADM). Tehran: Negahe danesh.

Bao, J., Bian, Z., Yu, Z., Phanphichit, T., Wang, G. and Zhou, Y., 2021, August. A Hybrid Approach to Risk Analysis for Critical Failures of Machinery Spaces on Unmanned Ships by Fuzzy AHP. In International Conference on Neural Computing for Advanced Applications (pp. 273–287). Springer, Singapore.

Bastan, M. and Shakouri, H., 2018. A system dynamics model for policy evaluation of energy dependency. In The 2nd IEOM European International Conference on Industrial Engineering and Operations Management.

Brouwer, P., van der Werf, A., Schluepmann, H., Reichart, G.J. and Nierop, K.G., 2016. Lipid yield and composition of Azolla filiculoides and the implications for biodiesel production. BioEnergy Research, 9(1), 369–377.

Buckley, JJ., Fuzzy hierarchical analysis. Fuzzy Sets and Systems 1985; 17:233–247.

Fallah, M., Pirali Zefrehei, A. and Hedayati, S.A., 2020. Determination of Potential Zones of Azolla filiculoides in Anzali International Wetland using GIS Technique. Health and Development Journal, 9(1), 29–42.

Firouzi, S., Nikkhah, A. and Aminpanah, H., 2018. Resource use efficiency of rice production upon single cropping and ratooning agro-systems in terms of bioethanol feedstock production. Energy, 150, pp.694-701.

Hassanzadeh, M., Zarkami, R. and Sadeghi, R., 2021. Uptake and accumulation of heavy metals by water body and Azolla filiculoides in the Anzali wetland. Applied Water Science, 11(6), 1–8.

Ho, W., He, T., Lee, C.K.M., and Emrouznejad, A., 2012. Strategic logistics outsourcing: An integrated QFD and fuzzy AHP approach. Expert Systems with Applications, 39(12), 10841–10850.

Hsieh, T.Y., Lu, S.T. and Tzeng, G.H., 2004. Fuzzy MCDM approach for planning and design tenders selection in public office buildings. International Journal of Project Management, 22(7), 573–584.

Jafari, N., 2009. Ecological integrity of wetland, their functions and sustainable use. Journal of Ecology and the Natural Environment, 1(3), 045–054.

Kaganski, s., Majak, J., & Karjust, K. (2018). Fuzzy AHP as a tool for prioritization of key performance indicators. Procedia CIRP.

Kamaruddin, N.A., Yusuf, N.M., Ishak, M.F. and Kamarudin, M.S., 2019. Study on Chemical Composition of Azolla filiculoides and Hydrilla verticillata. Journal of Agrobiotechnology, 10(1S), 68–74.

Laarhoven PJM, Pedrycz W. A fuzzy extension of Saaty ‘s priority theory. Fuzzy Sets Syst 1984;11(3):229 41.

Lotfi, M., Asgharizadeh, E., Hisam Omar, A., Hosseinzadeh, M. and Amoozad Mahdiraji, H., 2021. Measuring staff satisfaction in transportation system using AHP method under uncertainty. International Journal of Uncertainty, Fuzziness and Knowledge-Based Systems, 29(06), 875–889.

Maceika, A., Bugajev, A., Šostak, O.R. and Vilutiené, T., 2021. Decision tree and AHP methods application for projects assessment: a case study. Sustainability, 13(10), p.5502.

Madani, S. and Ravanbakhsh, M., 2018. A review of the status of invasive plant, case study: water hyacinth Distribution in Guilan Province. Human & Environment, 16(1), 129–138.

Maswada, H.F., El-Razek, A., Usama, A., El-Sheshtawy, A.N.A. and Mazrou, Y.S., 2021. Effect of Azolla filiculoides on growth, physiological and yield attributes of maize grown under water and nitrogen deficiencies. Journal of Plant Growth Regulation, 40(2), pp.558–573.

Mohamed, M.A., Elnemir, S.E., El-Mounem, A. and Abo El-Maati, S.M., 2018. Azolla fern as untraditional resource of protein. Zagazig Journal of Agricultural Research, 45(4), 1345–1355.

Nikkhah, A., 2018. Life cycle assessment of the agricultural sector in Iran (2007–2014). Environmental Progress & Sustainable Energy, 37(5), pp.1750–1757.

Pereira, A.L., 2017. The unique symbiotic system between a fern and a cyanobacterium, Azolla-Anabaena Azolla: Their potential as biofertilizer, feed, and remediation. In Symbiosis. IntechOpen.

Rajabi, M., Sardroud, J.M. and Kheyroddin, A., 2021. Green standard model using machine learning: identifying threats and opportunities facing the implementation of green building in Iran. Environmental Science and Pollution Research, 28(44), pp.62796–62808.

Razavipour, T., Moghaddam, S.S., Doaei, S., Noorhosseini, S.A. and Damalas, C.A., 2018. Azolla (Azolla filiculoides) compost improves grain yield of rice (oryza sativa L.) under different irrigation regimes. Agricultural Water Management, 209, pp.1–10.

Roy, D. C., M. C. Pakhira, and S. Bera. A review on biology, cultivation and utilization of Azolla. Adv Life Sci 5, no. 1 (2016): 11–15.

Saaty, T.L., (1980) The Analytic Hierarchy Process, McGraw-Hill, New York, USA.

Sadeghi, R., Zarkami, R. and Van Damme, P., 2014. Modelling habitat preference of an alien aquatic fern, Azolla filiculoides (Lam.), in Anzali wetland (Iran) using data-driven methods. Ecological modelling, 284, 1–9.

Sadeghi, R., Zarkami, R., Sabetraftar, K. and Van Damme, P., 2012. Application of classification trees to model the distribution pattern of a new exotic species Azolla filiculoides (Lam.) at Selkeh Wildlife Refuge, Anzali wetland, Iran. Ecological Modelling, 243, 8–17.

Salehzadeh, A., Naeemi, A.S. and Arasteh, A., 2014. Biodiesel Production from Azolla filiculoides (water fern). Tropical Journal of Pharmaceutical Research, 13(6), pp.957–960.

Singh, P.K., Panigrahi, B.C. and Satapathy, K.B., 1981. Comparative efficiency of Azolla, blue-green algae and other organic manures in relation to N and P availability in a flooded rice soil. Plant and Soil, 62(1), 35–44.

Thapa, P. and Poundel, K., 2021. Azolla: Potential biofertilizer for increasing rice productivity and government policy for implementation. J. Wastes Biomass Manag, 3, 62–68.

Vatandoost, M., Naghipour, D., Omidi, S. and Ashrafi, S.D., 2018. Survey and mapping of heavy metals in groundwater resources around the region of the Anzali International Wetland; a dataset. Data in Brief, 18, 463–469.

